# A systematically optimized awake mouse fMRI paradigm

**DOI:** 10.1101/2022.11.16.516376

**Authors:** Wenjing Xu, Mengchao Pei, Kaiwei Zhang, Chuanjun Tong, Binshi Bo, Jianfeng Feng, Xiao-Yong Zhang, Zhifeng Liang

## Abstract

Functional magnetic resonance imaging (fMRI) has been increasingly utilized in mice. Due to the non-negligible effects of anesthetics on mouse fMRI, it is becoming more common to perform fMRI in the awake mice. However, high stress level and head motion in awake mouse fMRI remain to be fully addressed, which limits its practical applications. Therefore, here we presented a systematically optimized awake mouse fMRI paradigm as a practical and open-source solution. First, we designed a soundproof habituation chamber in which multiple mice can be habituated simultaneously and independently. Then, combining corticosterone, body weight and behavioral measurements, we systematically evaluated the potential factors that may contribute to animals’ stress level for awake imaging. Among many factors, we found that the restraining setup allowing forelimbs freely moving and head tilted at 30-degree was optimal for minimizing stress level. Importantly, we implemented multiband simultaneous multi-slice imaging to enable ultrafast fMRI acquisition in awake mice. Compared to conventional single-band EPI, faster acquisition enabled by multiband imaging were more robust to head motion and yielded higher statistical power. Thus, more robust resting-state functional connectivity was detected using multiband acquisition in awake mouse fMRI, compared to conventional single-band acquisition. In conclusion, we presented an awake mouse fMRI paradigm that is highly optimized in both awake mice habituation and fMRI acquisition, and such paradigm minimized animals’ stress level and provided more resistance to head motion and higher statistical power.

## 1. Introduction

Functional magnetic resonance imaging (fMRI) (Ogawa et al., 1990) is one of the most important neuroimaging tools for noninvasively studying brain activity. The mouse is probably the most widely used model organism in preclinical research. Mouse fMRI has attracted more attention in recent years. However, previous mouse fMRI studies often generated inconsistent results (Adamczak et al., 2010; Bosshard et al., 2010; Bukhari et al., 2018; Grandjean et al., 2014; Jonckers et al., 2014; Nallasamy and Tsao, 2011; Nasrallah et al., 2014; Reimann et al., 2018; Schlegel et al., 2015; Tsurugizawa and Yoshimaru, 2021; Wu et al., 2017; Xie et al., 2020). Although efforts have been made to develop and optimize anesthesia methods to achieve robust and specific activation in mouse fMRI (Grandjean et al., 2014; Hamilton et al., 2017; Petrinovic et al., 2016; Schlegel et al., 2015; Shim et al., 2018), anesthetics still have non-negligible effects on mouse fMRI both in neural activities and physiological conditions (Franceschini et al., 2010; Masamoto and Kanno, 2012; Pan et al., 2015; Shim et al., 2018; Petrinovic et al., 2016; Schlegel et al., 2018, 2015). Therefore, it has become increasingly common to perform mouse fMRI in the awake condition (Gao et al., 2017; Chen et al., 2020; Desai et al., 2011; Han et al., 2019; Harris et al., 2015; Madularu et al., 2017; Takata et al., 2018; Tong et al., 2019; Yoshida et al., 2016). However, the awake mouse fMRI still suffers from some unresolved issues, most notably the high stress level and large head motion.

In awake mouse fMRI, the mouse’s head, and sometimes its body, is tightly restrained to limit movement during imaging, which can be very stressful for the mouse. Although mice are usually habituated to MRI scanning environment including loud scanning noise and restraining, their stress level typically lacks monitoring. Some studies recorded physiological indicators of mice during habituation, including ECG, EMG, and the mount or weight of fecal (Madularu et al., 2017; Tsurugizawa et al., 2020; Yoshida et al., 2016; Gutierrez-Barragan et al., 2022; Harris et al., 2015). However, these indicators can’t accurately reflect the stress level of mice. Some studies measured cortisol or corticosterone concentration (Almeida et al., 2021; Gutierrez-Barragan et al., 2022; Tsurugizawa et al., 2020; Harris et al., 2015), but found that the stress level of mice did not return to baseline even with 25-day habituation. Thus, a systematic evaluation and optimization on stress level in awake mouse fMRI is much needed.

Furthermore, imaging techniques for awake mouse fMRI also require further development and optimization for better motion robustness and higher acquisition efficiency. In human fMRI studies, simultaneous multi-slice (SMS) imaging techniques have significantly increased the temporal resolution in fMRI (Feinberg and Yacoub, 2012; Moeller et al., 2010; Poser and Setsompop, 2018; Uǧurbil et al., 2013). And fMRI with shorter repetition time (TR) is more robust to head motion (Chen et al., 2020). Moreover, test statistics and temporal signal-to-noise ratio (tSNR) may benefit from increase of the effective sample size (Feinberg et al., 2010; Jahanian et al., 2019; Todd et al., 2016), leading to more significant activation or functional connectivity in task (Boyacioğlu et al., 2015; Chen and Glover, 2015; Todd et al., 2017, 2016) and resting-state fMRI (Feinberg et al., 2010; Preibisch et al., 2015; Smith et al., 2013). However, SMS techniques have not yet been widely applied in small animal fMRI (Lee et al., 2019), and it is unknown whether SMS imaging would be particularly beneficial for awake mouse fMRI.

To this end, in the current work, we provided a highly optimized awake mouse fMRI paradigm, by both systematically optimized the habituation paradigm for reducing stress level and the multiband (MB) EPI for better motion robustness and higher acquisition efficiency. First, we designed a soundproof habituation chamber in which six mice can be habituated simultaneously and independently. Then, we systematically evaluated potential factors that might contribute to animals’ stress level using corticosterone measurements, body weights and behavioral tests. Those factors included 1) restraining setup designs; 2) environment enrichment; 3) earplugs; 4) sound intensity and 5) fMRI scanning duration. Based on the habituation paradigm, we further implemented MB EPI method combining SMS and generalized auto-calibrating partially parallel acquisitions (GRAPPA) (Griswold et al., 2002), which exhibited improved performance both in head motion robustness and statistical power. In conclusion, we presented a highly optimized awake mouse fMRI paradigm, which is an open-source solution (https://github.com/ZhifengLiangLab/awake-mouse-fMRI.git) for easy implementation.

## 2. Methods

### 2.1. Animals

All animal experiments were approved by the Animal Care and Use Committee of the Institute of Neuroscience, Chinese Academy of Sciences, Shanghai, China. Male C57BL/6 mice, 5-week-old, were used in the current study. Mice were group housed (6/cage) under a 12-h light/dark cycle (light on from 7 a.m. to 7 p.m.) with food and water ad libitum.

### 2.2. Head Holder implantation

The surgical procedure was conducted as previously (Chen et al., 2020; Han et al., 2019) when mice were 6-week old. Briefly, animals were anesthetized with 5% isoflurane for induction and 1–2% for maintenance. The surgical procedures were under the standard aseptic condition. The scalp, related soft tissues above the skull and muscle covering the skull posterior to the lambda point were removed. Then the skull was cleaned and dried, and a layer of light-curing self-etch adhesive (3M ESPE Adper Easy One) was applied to cover the interparietal bone and the surrounding regions. After the adhesive was light cured, a thick layer of light-curing flowable resin was applied on the same location, then a custom-made restraining setup was placed on the interparietal bone, and the resin was cured completely by blue light. Other exposed regions of the skull were covered with a thick smooth layer of dental cement to prevent the skull from inflammation. After the surgery, mice were given 7 days for recovery.

### 2.3. Habituation setup

To mimic the fMRI scanning environment, we designed a habituation chamber consisting of six independent small habituation boxes (Fig. 1B), allowing mice to be acclimated individually. The habituation chamber and six habituation boxes were assembled from sound attenuating panels (thickened wooden planks with a cement sandwich in the middle). The sound attenuation of the habituation chamber was measured at −50 dB, i.e., when the MRI scanning noise inside the habituation box reaches 110 dB, the noise outside the habituation chamber can be reduced to about 60 dB.

**Figure 1.**
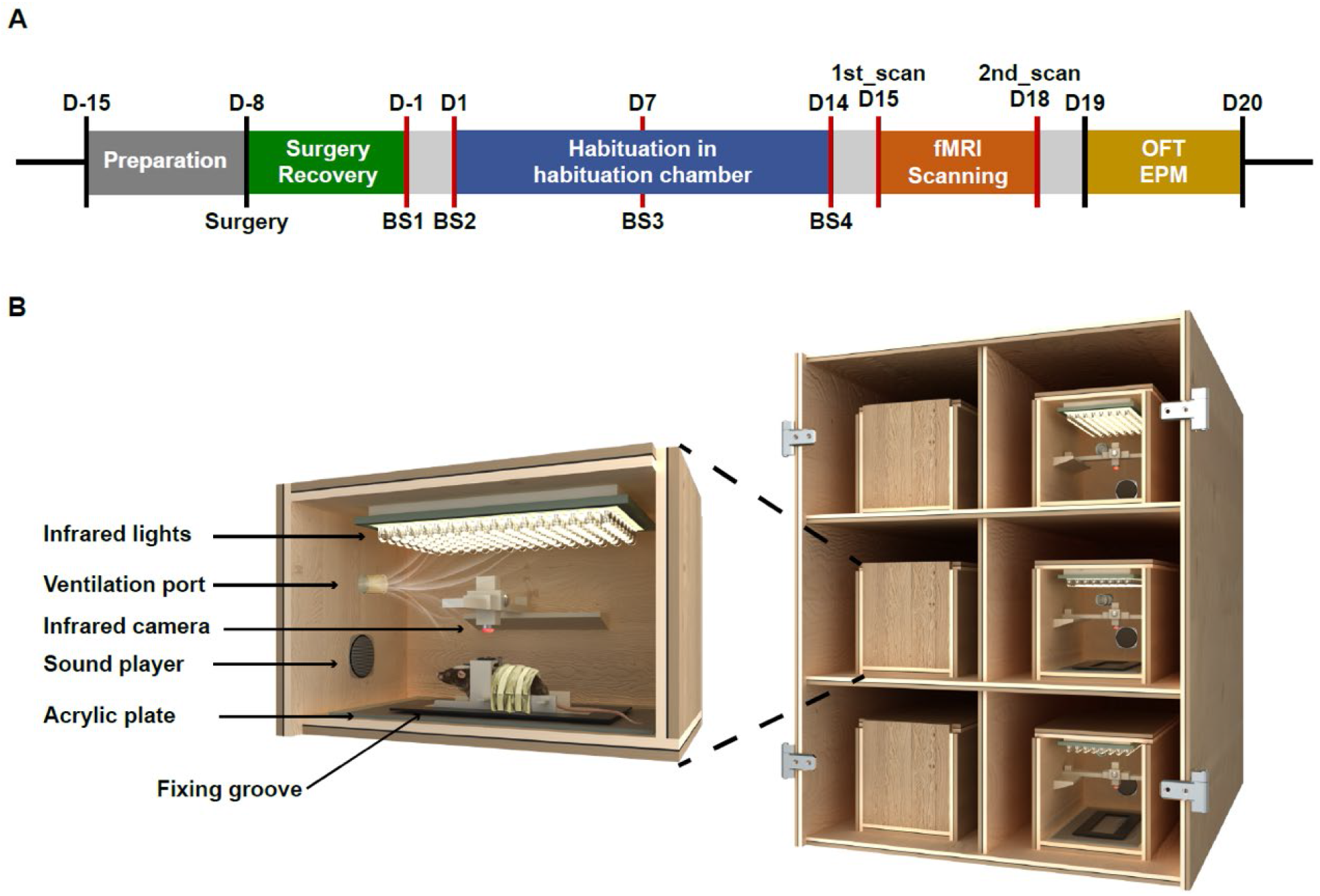
Overall study design (A) and schematic illustration of habituation setup (B).

Each individual habituation box includes five functional modules: 1) a sound player to playback the recorded MRI noise, with a manual switch to adjust the sound intensity; 2) a groove to fix the animal bed; 3) an infrared (IR) camera to monitor the mouse behavior with adjustable positions; 4) an IR light board to provide IR illumination; and 5) a ventilation hose to deliver fresh air to the habituation box. The design drawing of the habituation setup is publicly available (https://github.com/ZhifengLiangLab/awake-mouse-fMRI.git).

### 2.4. Optimization of animal restraining setup and habituation condition

To systemically optimize animal restraining setup (RS) and habituation condition, we specifically examined the effects of five factors: 1) RS designs (two factors: forelimb moving space and head tilting degree); 2) environment enrichment; 3) earplugs; 4) sound intensity; and 5) fMRI scanning duration.

Three types of RS designs were used and compared to examine the effects of forelimb moving space and head tilting degree. 1) The first type of RS design (RS1, Fig. 2A) was modified from those in our previous studies (Chen et al., 2020; Han et al., 2019). With this design, the mouse forelimbs were constrained under its head, and the head holder was fixed to the animal bed with screws so that the mouse head and forelimbs were tightly secured. 2) Based on RS1, the second design (RS2, Fig. 2B) was modified to raise the mouse forelimbs for 2.6 cm to allow more space for forelimb movement. 3) Based on RS2, a third design (RS3, Fig. 2C) was further modified so that the mouse head was tilted down 30 degrees, which is close to the mouse’s natural head position observed during quiet wakefulness and sleep (Yüzgeç et al., 2018). In addition, in all three designs, the mouse body was padded with sponges to make them as comfortable as possible while limiting their body movements. All 3D printing files for RS designs are openly accessed (https://github.com/ZhifengLiangLab/awake-mouse-fMRI.git).

**Figure 2.**
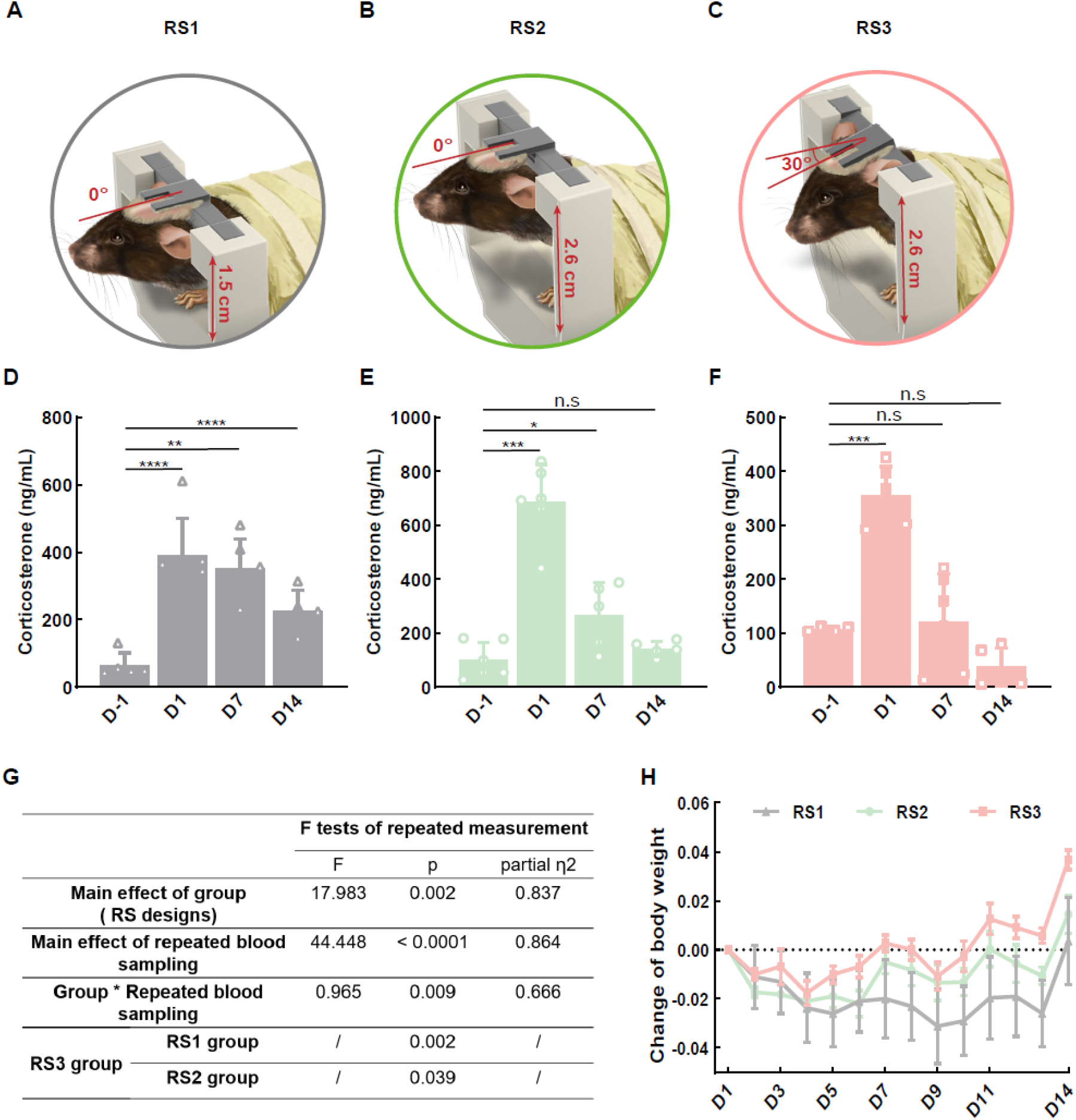
Effects of different restraining setups on corticosterone level and body weight during habituation in mice. (A) Restraining setup 1 (RS1): the mouse forelimbs were tightly constrained under its head. (B) Restraining setup 2 (RS2) modified from RS1: the mouse forelimbs were lifted up for 2.6 cm to allow more space for forelimb movement. (C) Restraining setup 3 (RS3) modified from RS2: the mouse head was tilted down 30 degrees. (D∼F) The corticosterone level measured at four time points in groups of RS1 (D), RS2 (E), and RS3 (F). (G) Repeated measurement ANOVA results. (H) Change of body weight (mean ± SEM) during 14-day habituation in groups of RS1, RS2, and RS3. (n = 6, * p <0.05, ** p < 0.01, *** p < 0.001, **** p < 0.0001)

To examine the effect of enriched environment, we housed the mice in the environment enriched cages (EE) consist of a transparent plastic box (460 mm × 315 mm × 200 mm), running wheels, wooden blocks with different shapes and colors, crawling tunnels with 5-cm internal diameter, and shredded paper scraps for nesting. Six mice were housed in each EE cage. The toys were changed once a week, and the colors and shapes were different from the last time to ensure the effect of EE. Mice were housed in EE cages from the first week until the end of the experiment, for a total of 5 weeks, which is sufficient for producing EE effects.

We also examined the effect of earplugs. First, the mouse ear canal was modeled using ABR impression material (SoundLink, Inc., Suzhou, China) to obtain the shape and size of its ear canal. Then, we resized the earplugs for human to fit the mouse ear canals. Before daily habituation, the modified earplugs were inserted into the mouse canals using forceps, and then the mixed impression material was injected into the mouse external ear canals to prevent the earplugs from falling off (Fig. S5E).

### 2.5. 14-day habituation procedure

After a week recovery from surgery, mice underwent habituation (once per day) for 14 days. Before daily habituation, mice were transported to the habituation room to acclimate to the environment for two hours, and the body weights of mice were measured and recorded. As shown in Table 1, the whole habituation period was divided into three stages: 1) The first 5 days, the habituation time gradually increased. Mice were fixed in animal beds and placed into habituation boxes without playing MRI scanning noise; 2) From day 6 to day 9, the MRI scanning noise intensity gradually increased; 3) From day 10 to day 14, mice were habituated to the maximum time (90 minutes) and maximum sound intensity (108 dB). In a real fMRI scanning, the sound intensity at the entrance of the magnet cavity was measured at about 103∼105 dB. As the sound would be presumably louder in the center of the magnet bore, a maximum sound level of 108 dB was used in the habituation.

### 2.6. Saphenous vein blood sampling

After comparing different blood sampling methods (Kim et al., 2018; Teilmann et al., 2014, 2012; Tsai et al., 2015), saphenous vein bleeding was chosen in the current study as it is less invasive and allows repeated blood collection with sufficient volume. To avoid the circadian rhythm effect on the corticosterone (Critchlow et al., 1963; Fahrenkrug et al., 2012; Gong et al., 2015; Thanos et al., 2009), all blood sampling (40 µl each time per mouse) was done between 11:00am and 2:00pm, immediately after habituation.

Specifically, the mouse was immobilized on the animal bed during blood sampling, and the hindlimb was shaved. The root of the hindlimb was pressed, and the skin was rubbed with 75% alcohol to dilate the blood vessels, and the saphenous vein was clearly visible. A 1 mL syringe needle, previously dipped in heparin solution (50 µl/mL), was used for puncture the saphenous vein. The blood sample was collected using heparin-coated glass capillary (Kimble Chase Life Science and Research Products, LLC, Vineland, NJ, USA) closed with the wax plate (Vitrex Medical A/S, Inc., Vasekær, Herlev, Denmark) at the end. Note that the entire blood collection procedure was completed within 2 minutes, minimizing the confounding stress related to the blood sampling itself.

The collected blood sample was stored in an ice box and then centrifuged. Capillaries with blood samples were placed in a hematocrit centrifuge (DM1244, Scilogex, LLC, CT, USA) and spun at 100 × 100 rpm for 10 min at room temperature to separate blood components. 15∼20 µl of the clear layer of plasma was transferred to an Eppendorf tube and placed in a −20 degrees freezer for further processing. After the entire experiment, all plasma samples were analyzed for enzyme-linked immunosorbent assay (ELISA).

### 2.7. Plasma Corticosterone Enzyme-Linked Immunosorbent Assay (ELISA)

Blood corticosterone level was measured with enzyme-linked immunosorbent assay (ELISA) following the manufacturer’s instructions. The DRG-4164 ELISA kit (DRG Instruments GmbH, Marburg, Germany) was used for its highest sensitivity (Kinn Rød et al., 2017), which is suitable for the current experiment with limited blood sample volume. Note that corticosterone was chosen rather than cortisol as a stress marker, as corticosterone has been reported to more directly respond to the chronic stress response (Gong et al., 2015). To ensure that the measured corticosterone concentration is within the measurement range of the kit, we used a gradient dilution rate in the experiment. The final samples after running all the procedures were read on a multi-mode microplate reader (FlexStation 3, Molecular Devices, California, USA). The assay was performed in duplicate. Custom-written scripts in MATLAB (MathWorks, Natick, MA) was used for plotting the standard curve and data extrapolation. Blood corticosterone level refers to the amount of corticosterone in plasma and is expressed in nanograms per milliliter (ng/ml).

### 2.8. Behavioral tests

It has been reported that chronic stress has a significant effect on mouse performance in the stress-sensitive behavioral paradigms (Chiba et al., 2012; Culig et al., 2017; Sadler and Bailey, 2016). Here, we used open field test (OFT) and elevated plus maze (EPM) to evaluate the effects of the habituation and imaging on the mouse’s behavior. In OFT, a white plywood box (40 × 40 cm2) was used, and its 20 × 20 cm2 center was defined as the central area. The room was illuminated at about 200 lux, and the light intensity of the central area and the surrounding area was the same. Each animal was placed in the center of this OFT box and each test lasted 10 min and was recorded with a video camera positioned above the center of the box. EPM consisted of two open arms (30 × 5 cm^2^), two closed arms (30 × 5 cm^2^) and a central area (5 × 5 cm^2^) made of yellow plastic, which was 50 cm above the ground. Each animal was first placed in the central area, and was recorded with a video camera placed above the center of the EPM for 5 min. Black curtains were hung around the EPM to produce low-light environments and ensure uniform light intensity for both open and closed arms.

Mice were transferred to the behavioral room two hours before the experiment and were tested individually. OFT test was performed on the first day and EPM on the other day to reduce the influence between two behavioral tests. To be consistent with corticosterone sampling time, all behavioral tests were performed between 11:00am and 2:00pm. All tests were performed by the same experimenter to avoid the influence of human operation. The software, EthoVision XT 13 (Noldus, Wageningen, the Netherlands) was used for analysis.

### 2.9. Multiband EPI sequence and reconstruction

The MB RF pulse was used for simultaneously multi-slice excitation. The waveform was designed as the sum of standard selective single-slice excitation pulse *A(t)* (e.g. in our fMRI study a Hanning windowed three-lobe sinc pulse was employed) with phase modulation of different offset frequencies, which can be expressed by Equation (1):

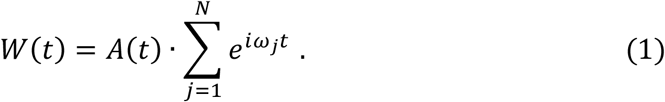

where the *W(t)* denotes the MB RF waveform and the *N* is MB factor (MBF), the *ωj* denotes the offset frequency of the jth slice.

The MB EPI pulse sequence is mainly made of two parts: 1) imaging scan and 2) reference scans:

1. The image scan is MB acquisition that the MB pulse *W(t)* is applied on the EPI pulse sequence (Fig. 4A) to simultaneously obtain MB signal, and it can run with (or without) combination of a phase encoding under-sampling technique, integrated parallel acquisition techniques (iPAT). Furthermore, the gradient blips train is applied on slice direction simultaneously with phase encoding blips to achieve CAIPIRINHA (Setsompop et al., 2012) – the approach of phase chopping different slice to shift away aliased slices in phase encoding direction to reduce parallel imaging g-factor by purpose.
2. The reference scans consist of three (four when iPAT was used) sequences:

i. Measurement of readout k trajectory which is obtained by subtracting the readout phase of two thin and close excited slice signals (Zhang et al., 1998).
ii. Phase correction scan by acquiring EPI signal without phase encoding blips (Fig. 4C). The phase inconsistency between EPI odd and even echoes is estimated by polynomial regressing (Bruder et al., 1992) and then finally canceled.
iii. Single-band (SB) calibration scan (Fig. 4B) for MB reconstruction that single-band pulse *A(t)* is used in EPI sequence to excite and acquire unaliased SB data which are matched with the MB imaging scan. The coefficients *λ* of MB reconstruction of different coils and slices are estimated by slice-GRAPPA algorithms (Setsompop et al., 2012), which are eventually used for reconstruct MB data of image scan by Equation (2):

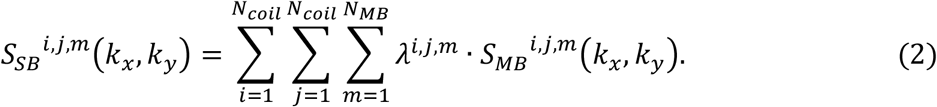

In which *i, j, m, N*_*coil*_ and *N*_*MB*_ are the index of coil of sampled data, index of coil of reconstructed data, index of slice, number of coils and MBF respectively. *S*_*MB*_ is the sampled MB data, and *S*_*SB*_ is the reconstructed unaliased single slice data.
iv. Auto-calibration signal (ACS) reference scan (Fig. 4D) for iPAT reconstruction that acquired full sampling but low-resolution data to estimate coefficients for phase encoding GRAPPA reconstruction (Griswold et al., 2002).

After image scan and reference scans finished, the MB imaging data were reconstructed step by step:

1. Re-gridding. The non-uniform Cartesian readout data (e.g. trapezoid ramp-sampling or sine function) was re-gridded by convolution k trajectory measured by reference scan i with a Kaise-Bessel convolution window;
2. Odd-even echo phase correction using regressions from reference scan ii;
3. Slice-GRAPPA reconstruction using the coefficients computed from reference scan iii;
4. Finally, if iPAT used, phase encoding GRAPPA reconstruction using the coefficients calibrated from reference scan iv.

### 2.10. MRI acquisition

MRI data were acquired with a Bruker BioSpec 9.4T scanner (Bruker BioSpin, Ettlingen, Germany), using 86 mm volume coil (Bruker) for transmission and 4 channel phased array cryogenic mouse head coil (Bruker) for receiving. To be compatible with mouse cryogenic head coil and avoid the signal-to-noise ratio (SNR) degradation due to the distance from the coil, we modified RS2 to RS4 (Fig. S7A). Resting-state fMRI data acquisition was performed immediately after the 14-day habituation. To compare the MB EPI with the conventional EPI, mice were randomly divided into two groups (12 mice in total, 6 mice for MBF =1 and R = 2 (GRAPPA-EPI), 6 mice for MBF = 2 and R = 2 (Biband-EPI)). Mice were scanned twice with one day apart from each scan. The mouse was first secured in the animal bed without any anesthesia, and placed inside the MRI scanner. T2 RARE anatomical images were first acquired (TE 11 ms, TR 3400 ms, field of view 16 × 16 mm^2^, matrix size 200 × 200, slice thickness 0.5 mm, number of slices 30, number of averages 2, RARE factor 8). After field-map-based local shimming within the mouse brain, single-shot GRAPPA-EPI or Biband-EPI was acquired. For GRAPPA-EPI, images were acquired with the following parameters: TE 15 ms, TR 1000 ms, flip angle = 52.6, bandwidth 300 kHz, field of view 20 × 20 mm^2^, matrix size 100 × 100, nominal slice thickness 0.48 mm (slice thickness 0.4 mm with a slice gap of 0.08 mm), number of slices 30, number of repetitions 300. For Biband-EPI, images were acquired with the following parameters: TE 15 ms, TR 500 ms, flip angle = 38.8, bandwidth 300 kHz, field of view 20 × 20 mm^2^, matrix size 100 × 100, nominal slice thickness 0.48 mm (slice thickness 0.4 mm with a slice gap of 0.08 mm), number of slices 30, number of repetitions 600 (Table. S1). 6 ∼ 7 EPIs were acquired for each mouse on each day.

### 2.11. fMRI data analysis

The data analysis flow was summarized in Fig. S1. All data were pre-processed using custom scripts in MATLAB (MathWorks, Natick, MA) and SPM12 (http://www.fil.ion.ucl.ac.uk/spm/). Raw images were first reconstructed and converted to NIFTI for further analysis. The mouse brain tissue was extracted manually using ITK-SNAP (http://www.itksnap.org/). The data of first 20 seconds in each run was discarded to ensure the magnetization to reach steady state. The images of each scan were slice-timing corrected (GRAPPA-EPI only) and realigned to the first volume of each scan. Then, framewise displacement (FD) was calculated using the modulus of the differential realignment parameters for each scan assuming a mouse brain radius of 5 mm (Power et al., 2012). Volumes with FD larger than 0.05 mm (1/4 voxel) and their neighboring volumes (−2∼4 for GRAPPA-EPI, −2∼6 for Biband-EPI) were discarded. We manually selected contaminated volumes in global signal (GS) according to the following criteria: 1) volumes with FD > 0.05 mm; 2) and their neighboring volumes between sharp increase and sharp decrease volumes (Figs. 5A, B). EPI images were registered to a mouse brain template (0.2 × 0.2 × 0.5 mm^3^) (http://www.imaging.org.au/AMBMC/Model) for group analysis, and then spatially smoothed using an isotropic Gaussian kernel with an in-plane full-width-half-maximum (FWHM) of 0.8 mm. To assess the impact of motion parameter regression, four regression-based motion corrections were compared:

1. 12 regressors: 6 head motion parameters and their corresponding first order derivatives (Satterthwaite et al., 2013);
2. 27 regressors: 12 motion related regressors and 15 non-brain tissue based principal components (PCs). The top 15 PCs of signals were calculated using tissues outside the brain, e.g., from the muscle of the jaw, and were used to model non-neural signal variations (Chuang et al., 2019).
3. 37 regressors: 12 motion related regressors and 25 PCs.
4. 47 regressors: 12 motion related regressors and 35 PCs.

The Pearson’s correlation coefficients between FD and GS were calculated to quantitatively measure the degree to which the head motion related signal was reduced using combinations of the above regressors. We found that 37 regressors are the minimal regressors to sufficiently reduce motion related signal for GRAPPA-EPI and Biband-EPI (Figs. S2A, B). After nuisance signal regression, band-pass filtering (0.01 ∼ 0.1 Hz) was subsequently performed.

### 2.12. Statistical analysis

One-way ANOVA was used to compare differences in intra-group corticosterone level measured at different time points. Two-way repeated measurement ANOVA was used to compare differences in inter-group corticosterone level (Fig. 2G) or changes in body weight measured at different time points, and the Greenhouse-Geisser correction for lack of sphericity was applied. Two-sample t-test was used to compare behavioral differences between the two groups. All data analysis was performed using GraphPad Prism 8 (GraphPad Software, CA, US).

To measure functional connectivity (FC) from the resting-state fMRI data, a seed-based correlation analysis was performed with seed time-courses defined as the averaged signals from each labeled brain area in the AMBMC atlas. The FC was assessed for bilateral 5 ROIs: visual cortex (Vis), the primary somatosensory cortex (SSp), hippocampus (Hip), thalamus (Tha), and caudate putamen (CPu). Pearson’s correlation coefficients between each voxel and seed time-courses were calculated to generate the FC map. Fisher’s z-transformation was used to convert correlation coefficients to z values. Second level analysis was then conducted on the individual z-maps using one-sample t-test (FWE corrected p < 0.05 and cluster size >=10 voxels). The difference in connectivity strength between Biband-EPI and GRAPPA-EPI was compared using two-sample t-test (uncorrected p < 0.05 and cluster size >=10 voxels).

## 3. Results

### 3.1. Overall study design and habituation chamber

The overall habituation and imaging procedure were summarized in Fig. 1A and Table 1. During this procedure, 4 blood sampling (BS1∼BS4, Fig. 1A) were conducted for measuring the corticosterone level before and during the 14-day habituation (Table 1). fMRI scanning was performed one day after the 14-day habituation. As chronic stress in rodents may result in decreased food intake leading to weight loss (Chiba et al., 2012; Janakiraman et al., 2017), we also recorded the weight of animals before daily habituation.

A custom designed sound attenuation habituation chamber (Fig. 1B) was used for all habituation, which includes six independent habituation boxes and thus allows six mice to be habituated simultaneously and independently. The habituation boxes include sound players to playback MRI noises, infrared (IR) cameras and IR illumination to allow behavioral monitoring (Fig. S3), and ventilation hoses to deliver fresh air. Therefore, the habituation chamber allows high-throughput and convenient habituation for awake mouse fMRI.

Given the time and cost for animal habituation in a real MRI scanner, most awake mouse fMRI studies habituated mice in habituation devices designed to simulate the MRI scanning environment. Different from the habituation devices in previous studies (Madularu et al., 2017; Yoshida et al., 2016), our newly designed habituation chamber can habituate multiple mice simultaneously and independently, largely avoiding the mutual interference between mice and the interference of the external environment on the mice.

### 3.2. Optimization and evaluation of stress level in awake mouse fMRI habituation

We systematically evaluated potential factors that may contribute to animals’ stress level using corticosterone measurements, body weights and behavioral tests. Those factors include 1) restraining setup (RS) designs (forelimbs moving space and the head tilting degree, Fig. 2); 2) environment enrichment (Fig. S4); 3) earplugs (Fig. S5); 4) sound intensity (Figs. S6A, B) and 5) fMRI scanning duration (Figs. S6C, D). Three types of RS designs were described in details in Methods.

First, during the 14-day habituation, mice exhibited a general trend of initial corticosterone increase and then decrease over time (Figs. 2D-F), regardless the three RS designs (Figs. 2A-C). The corticosterone level of RS1 group (Fig. 2A) did not reduce to baseline (Fig. 2D), but the other two groups (RS2, Fig. 2B and RS3, Fig. 2C) returned to baseline at day 14 (Fig. 2E) and day 7 (Fig. 2F), respectively. The repeated measurement ANOVA (Fig. 2G) showed a significant main effect of RS designs (p = 0.002), and between group analysis showed that RS3 group was significantly better than RS1 (p = 0.002) and RS2 (p = 0.039). Body weight is also frequently used to evaluate stress level. The increase of weight was highest in RS3 group and lowest in RS1 group (Fig. 2H), although the group main factor was not significant (p = 0.3036). These results showed that the RS designs significantly influenced the animals’ stress level in habituation and the RS3 (freely moving forelimbs and 30-degree tilted heads) was the optimal design for reducing stress level.

It was reported that enriched environment (EE) can effectively reduce the negative effect of stress (Pierce et al., 2018; Sano et al., 2019; Sztainberg and Chen, 2010). Therefore, we also tested the effects of EE on stress level of habituation. In the two groups of mice testing for the effects of EE, the restraining design of RS3 was used. One group was housed in EE cage (EE group) and the other group was housed in standard cage (control group). Compared to control group, the corticosterone level of EE group was significantly lower at baseline (p = 0.0007), but exhibited no significant difference during habituation (Fig. S4A). However, the body weight gain was significantly higher in EE group (p < 0.0001, Fig. S4B). The OFT and EPM tests showed that the total movement distance of EE group was significantly higher (Figs. S4C, D), indicating better motor ability in EE group. In addition, the time duration in center area in OFT was significantly higher in EE group (Fig. S4C). The results of body weight and behavioral tests suggest that enriched environment may reduce mice stress level and negative effects of head-fixed habituation, although the corticosterone level was not significantly different between two groups during habituation.

Loud MRI scanning noise may be a stressor for awake mice fMRI and thus we examined the effects of earplugs and sound intensity. Surprisingly, the corticosterone level in earplug group increased consistently over the course of the habituation, which was significantly worse than the control group (Fig. S5A). The group main factor in weight change was not significant (p = 0.1957, Fig. S5B). The total movement distance and time duration in center area in OFT were significantly lower in earplug group (Fig. S5C), while there was no significant difference between two groups in EPM (Fig. S5D). Therefore, earplugs significantly increased the stress level and induced stress-related behavioral phenotypes. Next, we examined whether sound intensity had any significant effect on mice stress level. Two groups of mice were habituated under 100 dB and 110 dB, respectively, and the stress level between two groups during 14-day habituation showed no significant difference and returned to baseline at day14 (Fig. S6A). The change of body weight in the two groups was similar (p = 0.6683, Fig. S6B). These results suggest that the sound intensity difference may not be a main stress factor, at least for the difference between 100 dB and 110 dB.

We also examined the effect of fMRI scanning duration on mice stress level. Two groups of mice were imaged for 30 min and 60 min, respectively. As shown in Fig. S6C, there was no significant difference in stress level between the two groups during 14-day habituation. However, in repeated fMRI scanning, mice underwent longer imaging time (60 min) had significantly higher stress level than mice received shorter imaging time (30 min) (Fig. S6C). The change of body weight in the two groups during habituation was similar (p = 0.5473, Fig. S6D). Therefore, reducing the imaging time may be beneficial for reducing stress level.

Further, we examined the effects of 14-day habituation and repeated MRI scanning on mouse behavior. 12 mice secured with RS4 (Fig. S7A) were randomly and equally divided into two groups (Biband-EPI and GRAPPA-EPI) and were scanned two times within four days after 14-day habituation. Their corticosterone level returned to baseline at day 14 (Fig. 3A) and the change of body weight showed no significant difference between the two groups (p = 0.9244. Fig. 3B). OFT and EPM were performed on mice one day after repeated imaging. Compared to a control group in which 6 mice secured with RS4 but without habituation and fMRI scanning, there was no significant behavioral difference between these groups in OFT (Fig. 3C) and EPM (Fig. 3D), suggesting that the 14-day habituation and repeated fMRI scanning did not induce stress-related behavioral phenotypes.

**Figure 3.**
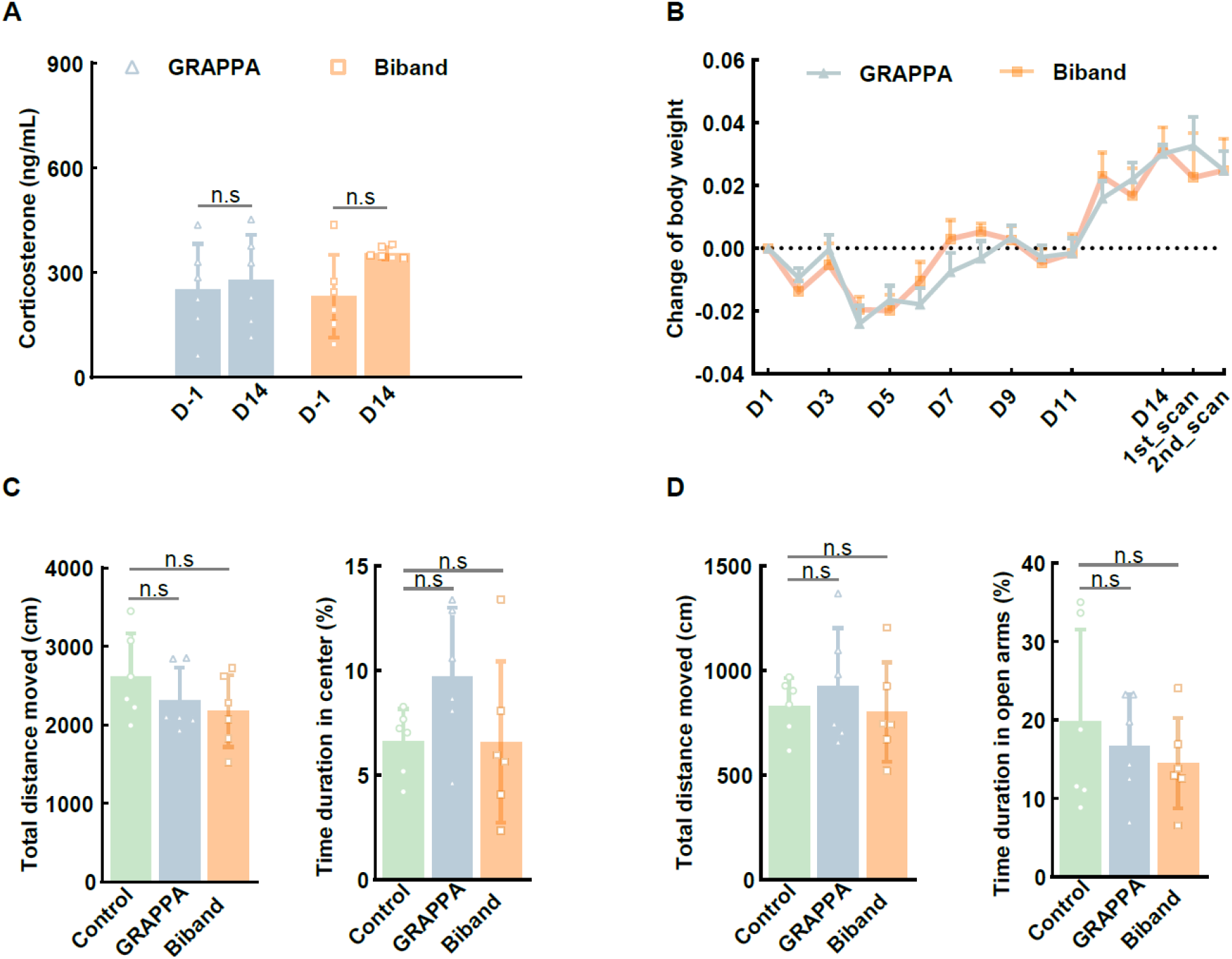
Effects of 14-day habituation and repeated MRI scanning on mice behavior. Corticosterone levels measured at Day-1 and Day14 in two groups. (B) Change of body weight (mean ± SEM) during habituation and repeated MRI scanning in two groups. (C, D) Open field test (OFT) and elevated plus maze (EPM) results in GRAPPA, Biband, and control groups, respectively. (n = 6)

### 3.3. Development and evaluation of multiband-EPI for awake mouse fMRI

To evaluate the performance of multiband-EPI in awake mouse fMRI, we developed a Biband-EPI (MBF = 2) combining with iPAT (acceleration factor R = 2) sequence, which is described in details in Methods (Figs. 4A-D). In Biband-EPI, a 2-band RF pulse was used to excite slices that were 7.2 mm apart in the slice-select direction. The blipped gradient increased the phase of simultaneously excited slices by π so that effectively shifts the acquired slices FOV/2 apart from each other. GRAPPA-EPI uses iPAT but no MB acceleration (MBF = 1 and R = 2). The Biband-EPI sequence and reconstruction software are openly accessed (https://github.com/ZhifengLiangLab/awake-mouse-fMRI.git).

**Figure 4.**
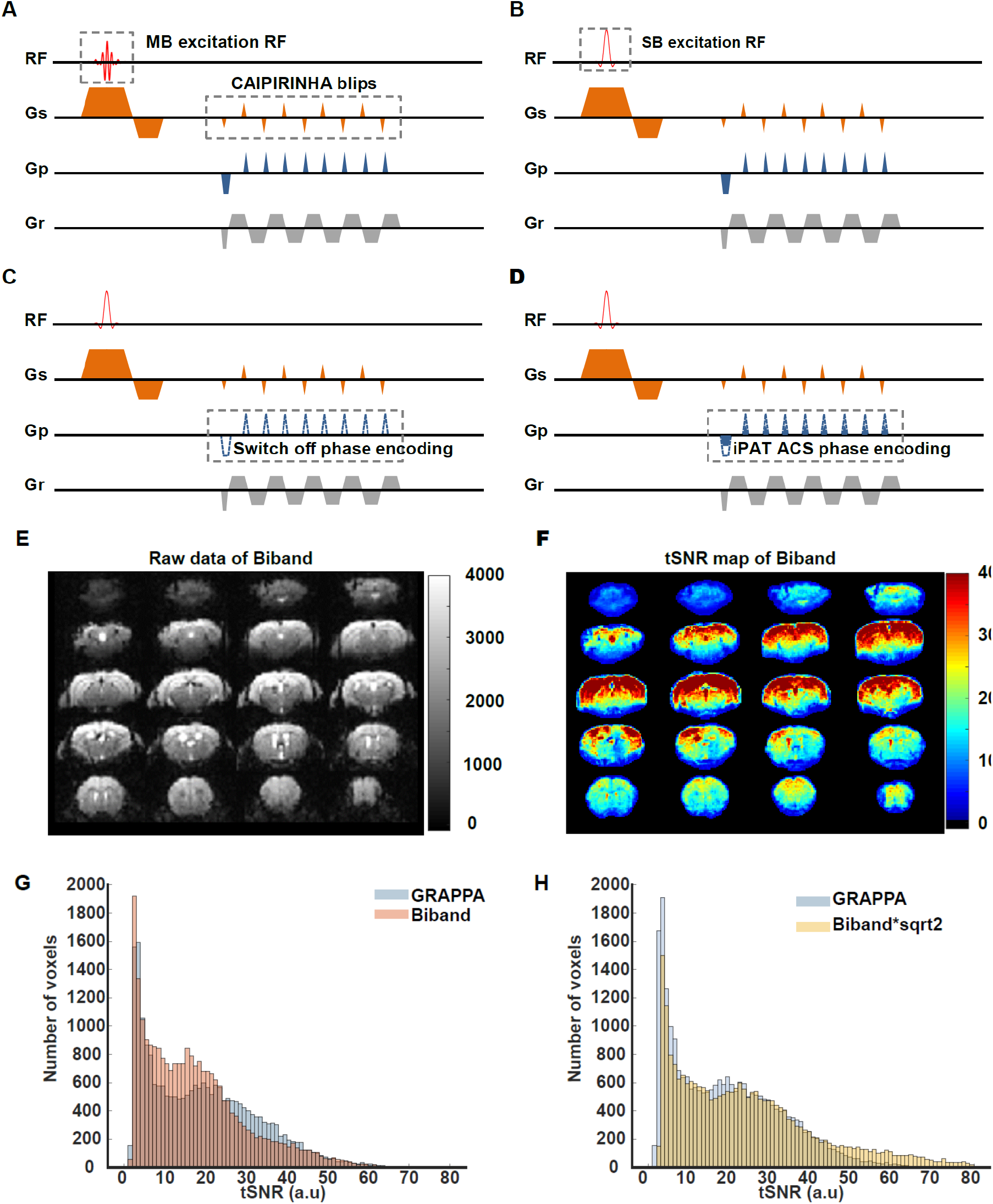
The pulse sequence of accelerated EPI with the combination of multiband (MB) and integrated parallel acquisition techniques (iPAT) acquisition. (A) Example of EPI sequence with an acceleration factor of 2×2 (MB factor (MBF) = 2; iPAT acceleration factor R = 2). CAIPIRINHA gradient was blipped on slice axis. (B) Single band (SB) reference scan using SB excitation RF pulse to obtain signal for calibrating coil coefficients of MB reconstruction. (C) None-phase encoding reference scan for EPI odd-even echo inconsistencies phase correction. (D) Auto-calibration signal (ACS) reference scan used to collect signal without GRAPPA phase encoding acceleration for estimating coil coefficients of GRAPPA reconstruction. (E) Raw Biband EPI data. (F) Temporal signal-to-noise ratio (tSNR) map of Biband-EPI. (G) tSNR histogram of Biband-EPI and GRAPPA-EPI. (H) Statistical power of Biband-EPI and GRAPPA-EPI.

Example slices of raw EPI images and the corresponding tSNR maps of Biband-EPI and GRAPPA-EPI from one representative animal were shown in Figs. 4E, F and Figs. S7B, C, respectively. Globally, since the TR of Biband-EPI was half of that in GRAPPA-EPI (500 ms vs. 1000 ms), the tSNR of Biband-EPI was slightly lower than the GRAPPA-EPI (Fig. 4G), particularly in subcortical and cerebral areas, but comparable in cortex and hippocampus (Fig. 4F and Fig. S7C). For fMRI analysis, the statistical power is related to tSNR and degree of freedom (DoF) and can be calculated by tSNR×√N, where N is the MBF (Chen et al., 2019; Smith et al., 2013). Therefore, as expected, the statistical power in Biband-EPI, with more data points acquired, was about 1.4 times higher than GRAPPA-EPI (Fig. 4H). Overall, it is clear that faster acquisition achieved by Biband-EPI is beneficial for awake mouse fMRI, just as in human fMRI.

Next, we evaluated whether faster acquisition speed by Biband-EPI would afford better robustness to head motion, which can be particularly problematic in awake animal imaging. The FD of four representative Biband-EPI and GRAPPA-EPI scans were small at most time points, but had some motion spikes (FD > 0.05 mm), which were also evident in GS time series (Figs. 5A, B). And motion spikes lasted longer in GS than in FD. As shown in Fig. 5C, head motion spikes contaminated about 6 - 8 frames in GS, including 2 frames before and 4 frames after in GRAPPA-EPI or 2 frames before and 6 frames after in Biband-EPI. Because Biband-EPI acquires 2-fold as many frames as GRAPPA-EPI, the proportion of contaminated frames in Biband-EPI is less. The number of frames with FD larger than 0.05 mm was significantly less in Biband-EPI (p < 0.001, Fig. 5D). In order to eliminate frames contaminated by motion spikes, we scrubbed these “bad” points from data. Compared to GRAPPA-EP, the number of scrubbed frames was significantly less in Biband-EPI (p < 0.01, Fig. 5E); and the mean FD was significantly lower in Biband-EPI before (p < 0.0001) and after (p < 0.001) scrubbing (Fig. 5F). Moreover, the correlation between FD and GS was also significantly lower in Biband-EPI before (p < 0.0001) and after (p = 0.0003) 37 nuisance signal regression (Figs. S2A, B). These results suggested that Biband-EPI was more robust to head motion.

**Figure 5.**
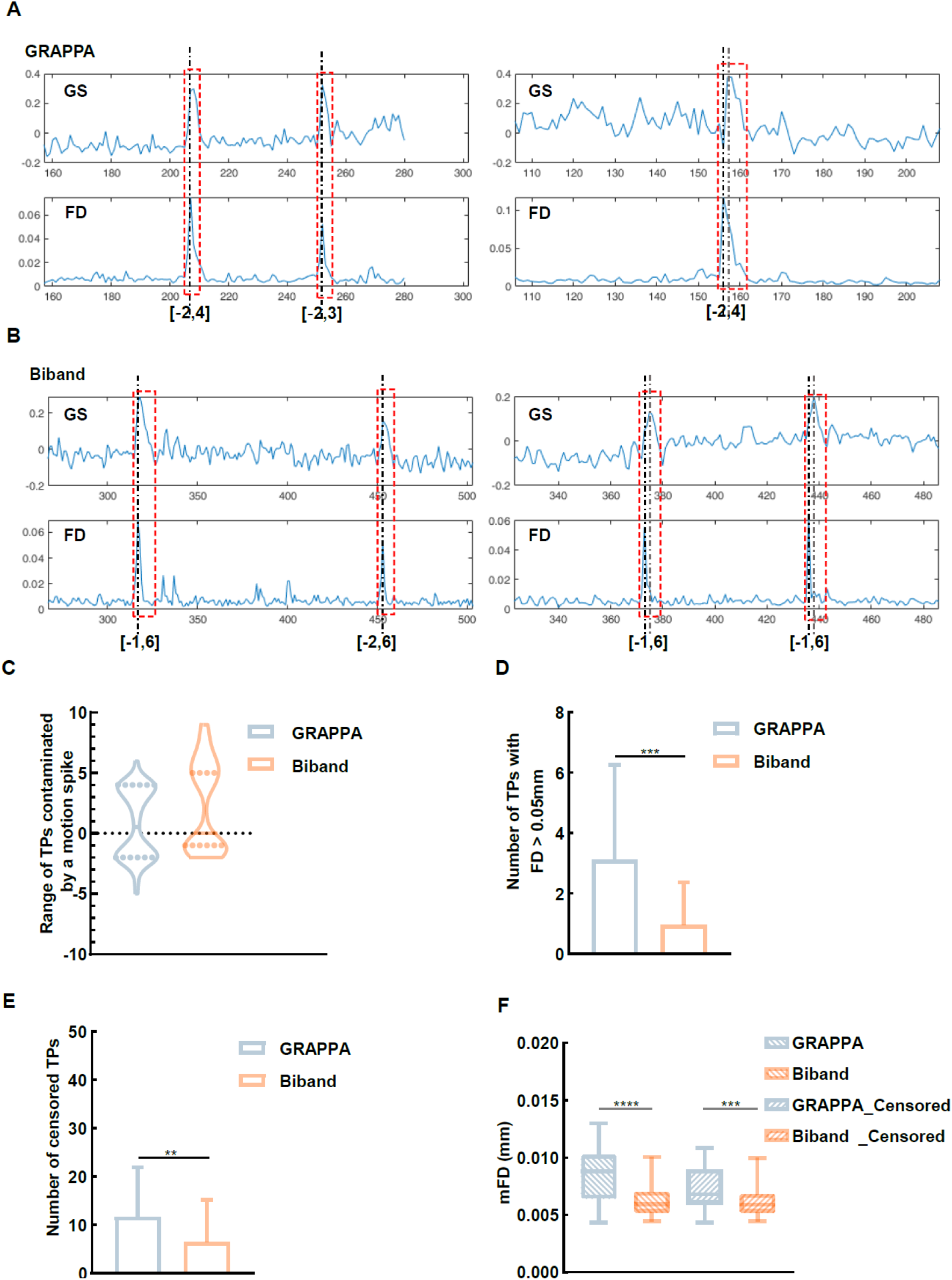
Comparison of head motion in GRAPPA-EPI and Biband-EPI data. (A, B) Framewise displacement (FD) and global signal (GS) of two fMRI scans from GRAPPA-EPI and Biband-EPI. (C) The range of time points (TPs) contaminated by a motion spike (FD > 0.05 mm) in GRAPPA-EPI and Biband-EPI. The mean contaminated TPs were (−1.7∼3.2) and (−1.3∼5.4) in GRAPPA-EPI and Biband-EPI, respectively. (D) The number of TPs with FD > 0.05 mm in GRAPPA-EPI and Biband-EPI. (E) The number of censored TPs in GRAPPA-EPI and Biband-EPI. (F) Comparison of mean FD (mFD) before and after censoring contaminated TPs in GRAPPA-EPI and Biband-EPI. (** p <0.01, *** p < 0.001, **** p< 0.0001)

Finally, we evaluated whether Biband-EPI would be beneficial for FC analysis in awake mice. As shown in Fig. 6, compared to GRAPPA-EPI, more robust resting-state FC was detected using Biband-EPI for 5 cortical and subcortical brain regions, including Vis, SSp, Hip, Tha, and CPu. For example, the t-values in SSp and Hip were significantly higher in Biband-EPI (Fig. 7), and correspondingly, higher statistical power leading to more spatial FC (Fig. 6). These results indicated that Biband-EPI was particularly beneficial for awake mice fMRI with greater robustness to head motion and higher statistical power.

**Figure 6.**
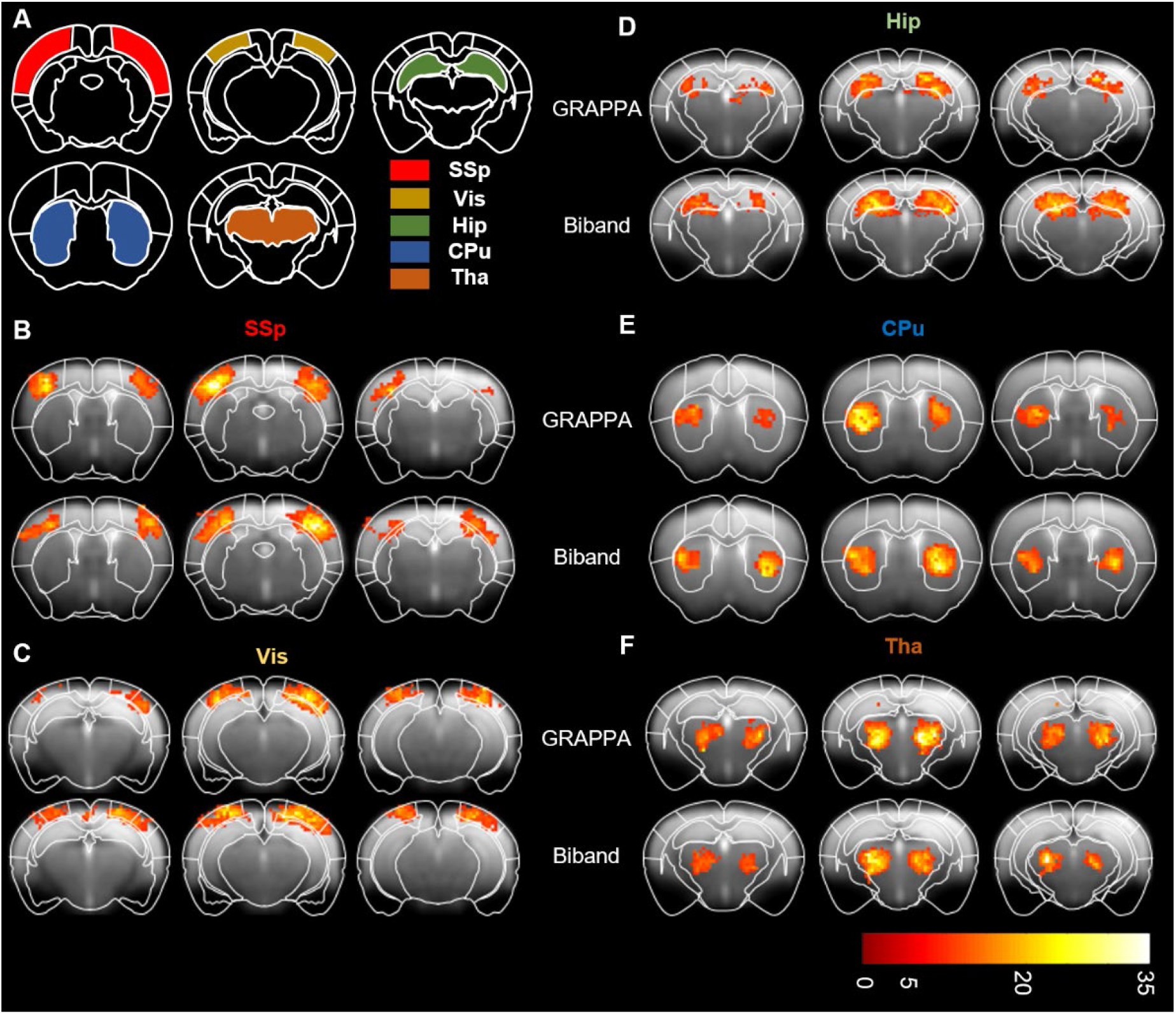
The resting-state functional connectivity (FC) of GRAPPA-EPI and Biband-EPI. (A) The region of interest (ROI) definition. SSp, primary somatosensory cortex; Vis, visual cortex; Hip, hippocampus; CPu, caudate putamen; Tha, thalamus. (B-F) The FC maps of those five ROIs, respectively. (one-sample t test with FWE correction, p < 0.05, voxel size >= 20, colorbar: T value)

**Figure 7.**
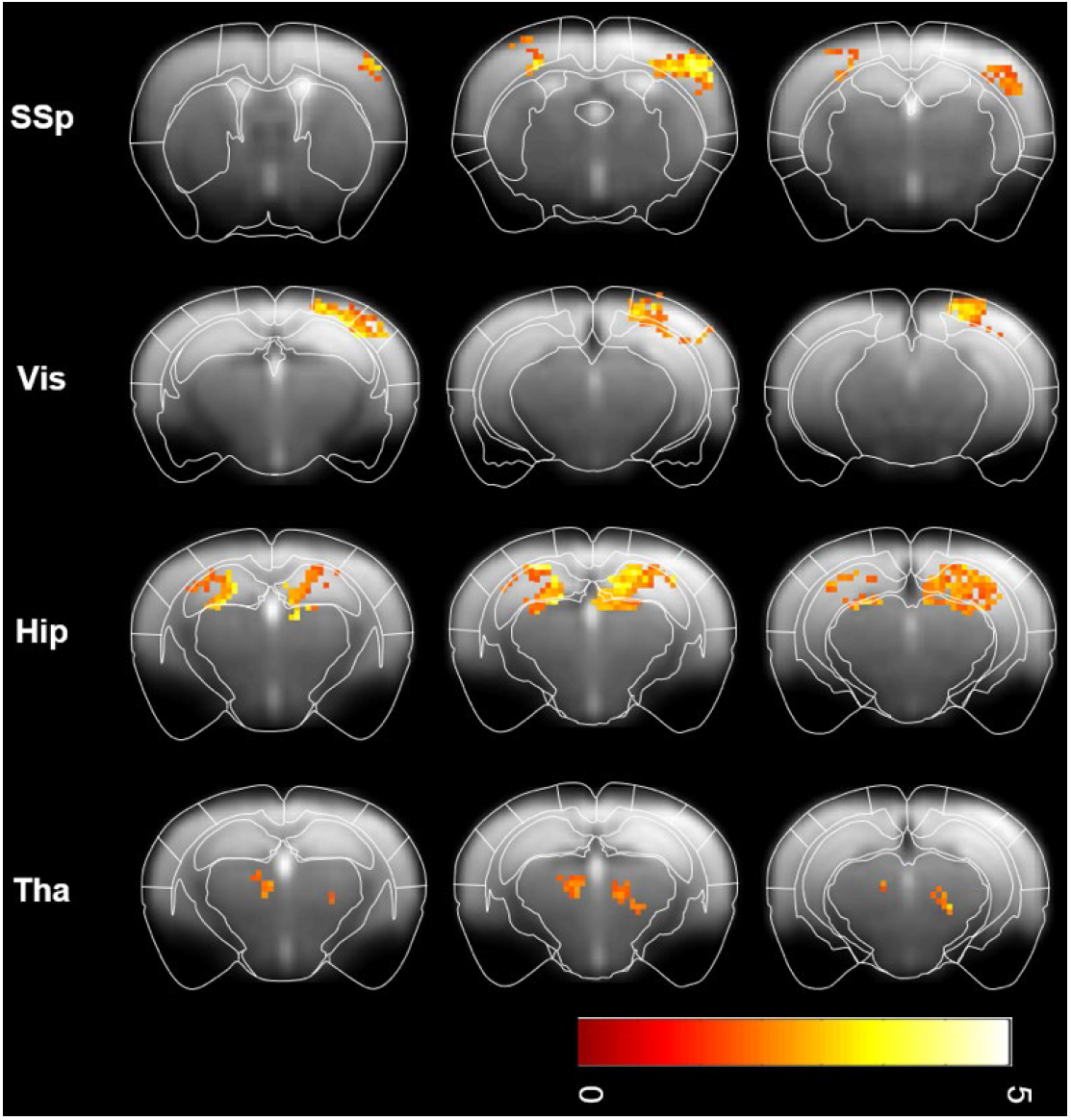
The comparison of resting-state FC between GRAPPA-EPI and Biband-EPI. The FCs in Biband-EPI were stronger than GRAPPA-EPI in SSp, Vis, Hip and Tha, but not in CPu. (two-sample t-test without correction p<0.05, voxel size >= 10, colorbar: T value)

## Discussion

In this work, we proposed an open-source and highly optimized awake mouse fMRI paradigm, with a systematically optimized habituation paradigm for reducing stress level and the multiband EPI for better motion robustness and higher acquisition efficiency.

Stress level is perhaps the biggest concern in awake animal fMRI. While it is not possible to completely eliminate the stress response, it is critical to evaluate and minimize the stress during awake imaging. Most current awake mouse fMRI studies lack stress level monitoring. Although some studies measured the respiratory rate, heart rate, and weight of fecal during habituation (Madularu et al., 2017; Tsurugizawa et al., 2020; Yoshida et al., 2016; Gutierrez-Barragan et al., 2022; Harris et al., 2015), these physiological indicators can’t represent the stress level directly. Some studies used cortisol or corticosterone concentration as the gold standard for measuring stress level (Almeida et al., 2021; Gutierrez-Barragan et al., 2022; Tsurugizawa et al., 2020; Harris et al., 2015), but most of these studies lack of intra-group longitudinal comparisons. In this study, we used plasma corticosterone as a gold indicator of stress level with longitudinal measurement to reflect the change of stress level accurately. While the corticosterone is known to be related to gender and circadian rhythm (Fahrenkrug et al., 2012; Gong et al., 2015; Thanos et al., 2009), only male mice were used in our study and all blood sampling was performed between 11:00am and 2:00pm. Besides, we incorporated behavioral tests and body weight monitoring. Thus, our work represented a comprehensive and effective approach to monitor stress level of awake mice.

Such comprehensive stress monitoring further enabled us to systemically evaluate potential stress factors with an ultimate goal of minimizing the stress level in awake mouse fMRI. As detailed in the method and results sections above, we made significant amount of efforts to evaluate various stress factors. Importantly, with the optimized restraining setup designs, mice stress level returned to baseline, which is a major improvement compared with previous studies (Gutierrez-Barragan et al., 2022; Tsurugizawa et al., 2020; Harris et al., 2015) using tight restraining similar to RS1. This indicates that tight restraining of the forelimbs is a major stressor for mice. We also found that tilting the head for 30 degrees was optimal, which was adopted from a study that found the head angle of mice was about 30 degrees during quiet wakefulness and sleep (Yüzgeç et al., 2018). A surprising finding was that earplugs made mice increasingly stressful over the habituation course and induced stress-related behavioral phenotypes. Therefore, earplugs may not be a good choice in awake fMRI to reduce stress level. Furthermore, we found that sound intensity may not be a main stressor (Figs. S6A, B), which is consistent with previous study (Almeida et al., 2021). This may attribute to the different hearing range and frequency sensitivity of the mouse. Unlike humans, mice have a wider hearing range (up to 100kHz) and are more sensitive to higher frequencies (around 10kHz). The recorded EPI noise is in a relatively low frequency range (Fig. S8). Another possible reason is that C57 mice are known to have hearing loss after 8 weeks (Parham, 1997; Walton et al., 2008). Therefore, for awake mouse fMRI, scanning noise may not be as critical as previously assumed.

Moreover, in the current study we provided the first practical SMS technique for high temporal resolution fMRI in awake mice, which is naturally applicable to other fMRI studies in awake or anesthetized animals. Compared to the only previous SMS EPI work in mouse fMRI (Lee et al., 2019), our Biband-EPI pulse sequence is more compatible with other phase encoding under-sampling techniques (e.g. iPAT, partial Fourier and zero filling) to further improve fMRI acceleration. And it also provides an option of reversed phase encoding scan for correcting field inhomogeneity distortion using TOPUP (Andersson et al., 2003; Graham et al., 2017). It does not require extensive additional data partly reconstructed by Bruker system (i.e., the data of each channel is re-gridded, phase corrected and separately saved as raw data). Instead, our program features the complete reconstruction, including re-gridding, phase correction, de-apodization, GRAPPA, etc., which requires original raw data only. Our offline reconstruction program which was implemented with Microsoft Visual Studio C++ on Windows platform is fast, robust and user-friendly. Thus, the compatibility and the reconstruction efficiency of our MB-EPI are greatly improved.

The better acquisition efficiency of MB-EPI is particularly useful for awake animal fMRI. First, shorter TR (with same spatial resolution and slice numbers) enabled by MB-EPI provides better robustness to head motion (Fig. 5), consistent with previous human studies (Smith et al., 2013; Uǧurbil et al., 2013). In addition, faster acquisition requires shorter imaging time, which may reduce animals’ stress level (Figs. S6C, D). The overall resting-state fMRI results in the current study clearly demonstrated increased sensitivity of FC in awake mice (Fig. 7). The MB-EPI parameters used in the current study mainly aimed to improve temporal resolution (TR 500ms vs. TR 1000ms in SB-EPI). If the temporal resolution remains the same to SB-EPI, better spatial resolution or spatial coverage (slice numbers) can be achieved in MB-EPI. Currently, MB-EPI is not commonly used for animal fMRI due to various reasons, e.g., lack of vendor-supplied sequences and reconstructions, low numbers of coil elements. Given the clear advantages demonstrated in the current study, we hope MB-EPI will be more widely used.

There were still some limitations in our study. First, we did not measure the corticosterone after MRI scanning but used behavioral tests to evaluate the effect of MRI scanning on the mouse’s stress level. The reason is mainly because the operation of removing mice from animal beds before blood sampling significantly influenced animal’s state so that the corticosterone cannot reflect animal stress level during scanning. In addition, multiple animals were imaged sequentially throughout the day, thus blood samples cannot be collected simultaneously to avoid the circadian rhythm effect on corticosterone level. Second, we did not evaluate the performance of Biband-EPI in task fMRI in awake mice, which will be performed in future studies. Third, we did not habituate mice in a real MRI scanner and did not compare the effects between habituation in MRI scanner and habituation chamber. In practice, the feasibility of habituation in MRI scanner is extremely limited because of high cost of time and scanner availability. Fourth, the physiological parameters, such as respiratory rate, heart rate and rectal temperature were not recorded during fMRI. Ideally, these parameters can provide references for us to evaluate the stress level of mice during imaging. Overall, future work is needed to further refine the awake mouse fMRI.

In conclusion, we provided a systematically optimized paradigm for awake mouse fMRI, with a habituation paradigm for minimizing stress level and an optimized MB-EPI sequence for high temporal fMRI in awake mouse fMRI. This paradigm provides a timely solution for the field of small animal fMRI, and is expected to further promote the applications of awake mouse fMRI in neuroscience.

## Acknowledgements

This study was supported by the National Natural Science Foundation of China (81873893, 82171903 and 82171899), National Science and Technology Innovation 2030 Major Program (2021ZD0200100 and 2021ZD0202200), Strategic Priority Research Program of Chinese Academy of Sciences (XDBS01030100), Lingang Laboratory (LG202104-02-06) and the Shanghai Municipal Science and Technology Major Project (2018SHZDZX01 and 2018SHZDZX05).

## Data availability

The whole awake mouse fMRI solution, including hardware setup and SMS sequence and reconstruction, is publicly available (https://github.com/ZhifengLiangLab/awake-mouse-fMRI.git).

