## Supplementary figures and images for "A systematically optimized awake mouse fMRI paradigm"

### Supplemental Figure 1

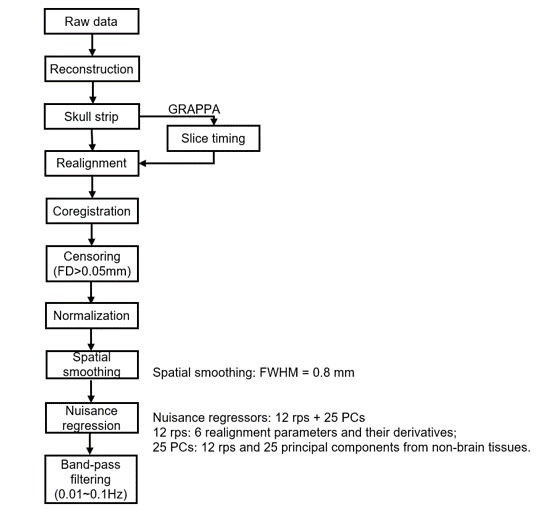

Figure S1. fMRI data preprocessing pipeline.
