## Supplemental Figure 2 for "A systematically optimized awake mouse fMRI paradigm"

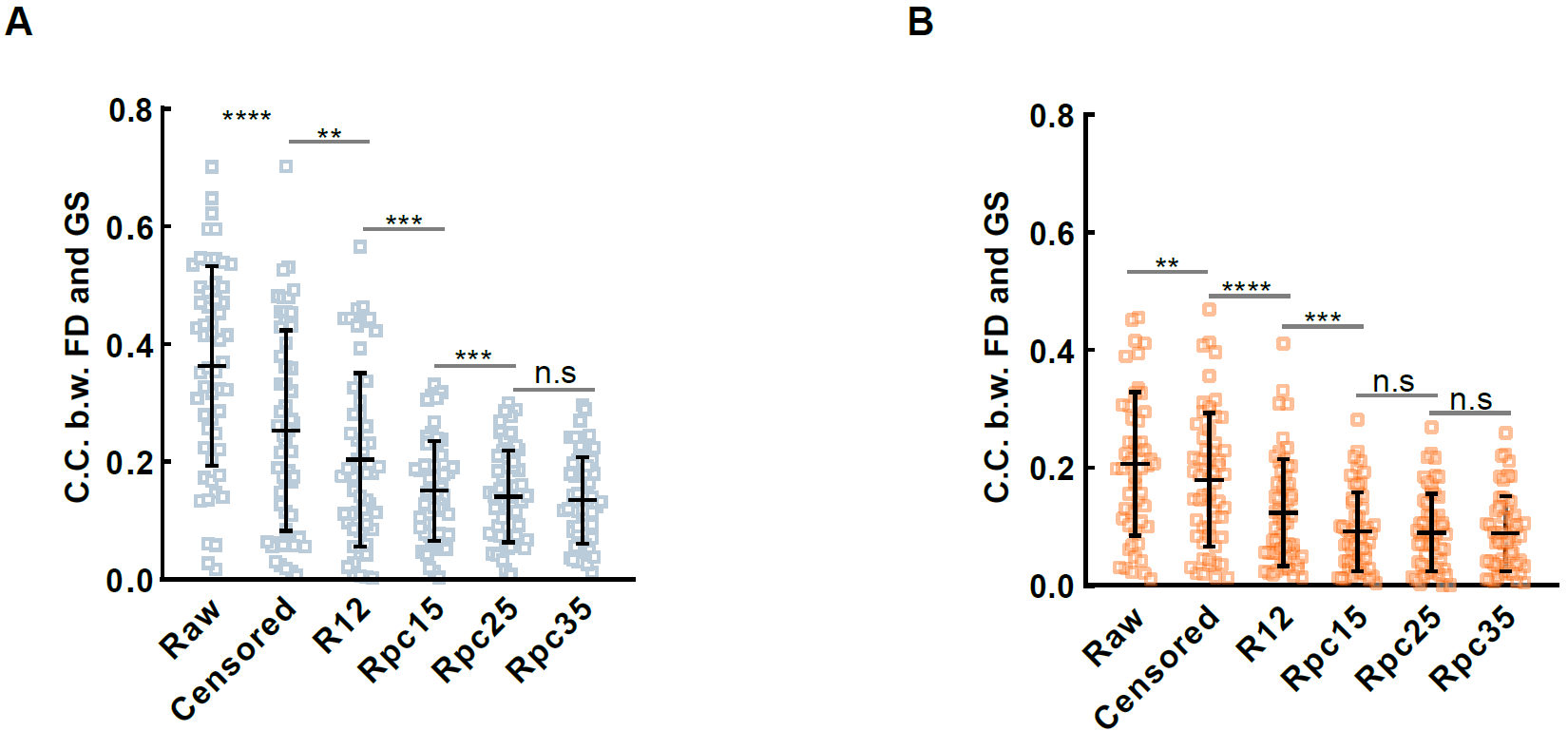

Figure S2. The Pearson’s correlation coefficients between FD and GS under different nuisance regressors in GRAPPA-EPI (A) and Biband-EPI (B). (** p<0.01, *** p < 0.001, **** p<0.0001)
