## Supplemental Figure 3 for "A systematically optimized awake mouse fMRI paradigm"

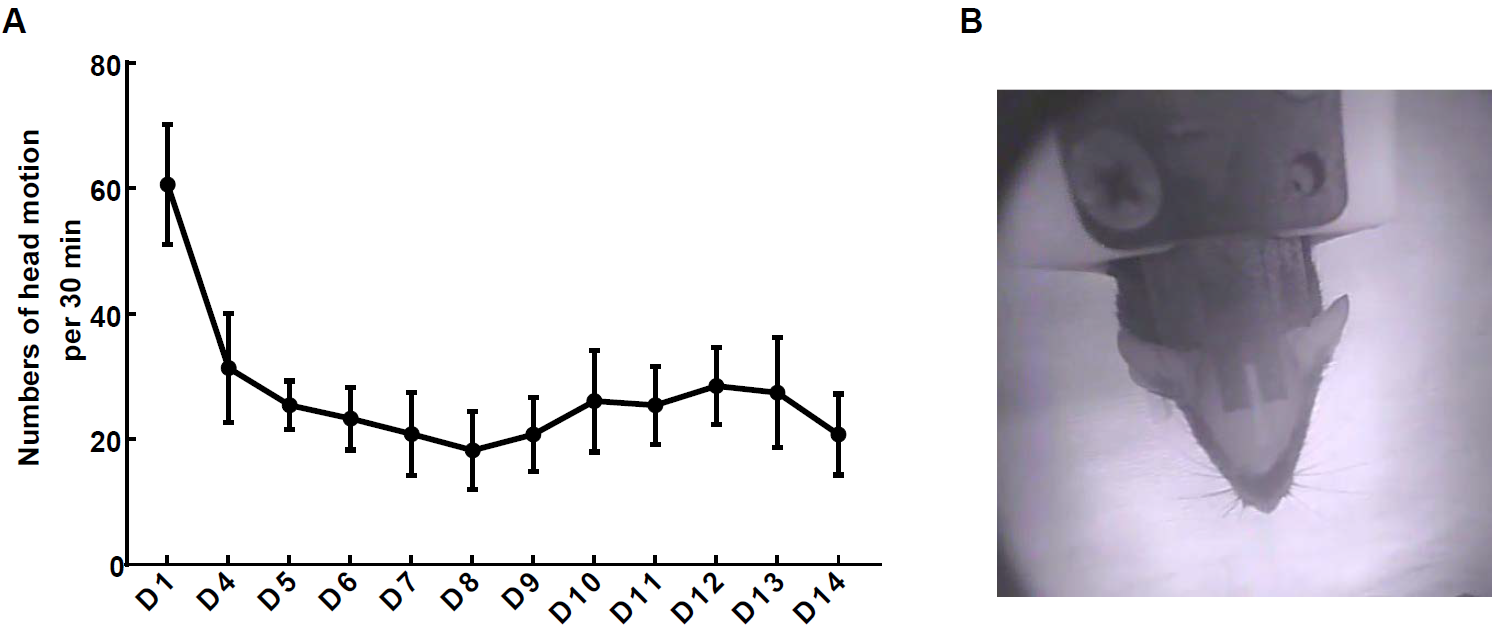

Figure S3. Numbers of mouse head motion (A) analyzed from the video recorded (B) during 14-day habituation (n=10). The numbers of motion reduced at first seven days and remained until the last day. Inter-frame difference method was used to extract the head motion.
