## Supplemental Figure 4 for "A systematically optimized awake mouse fMRI paradigm"

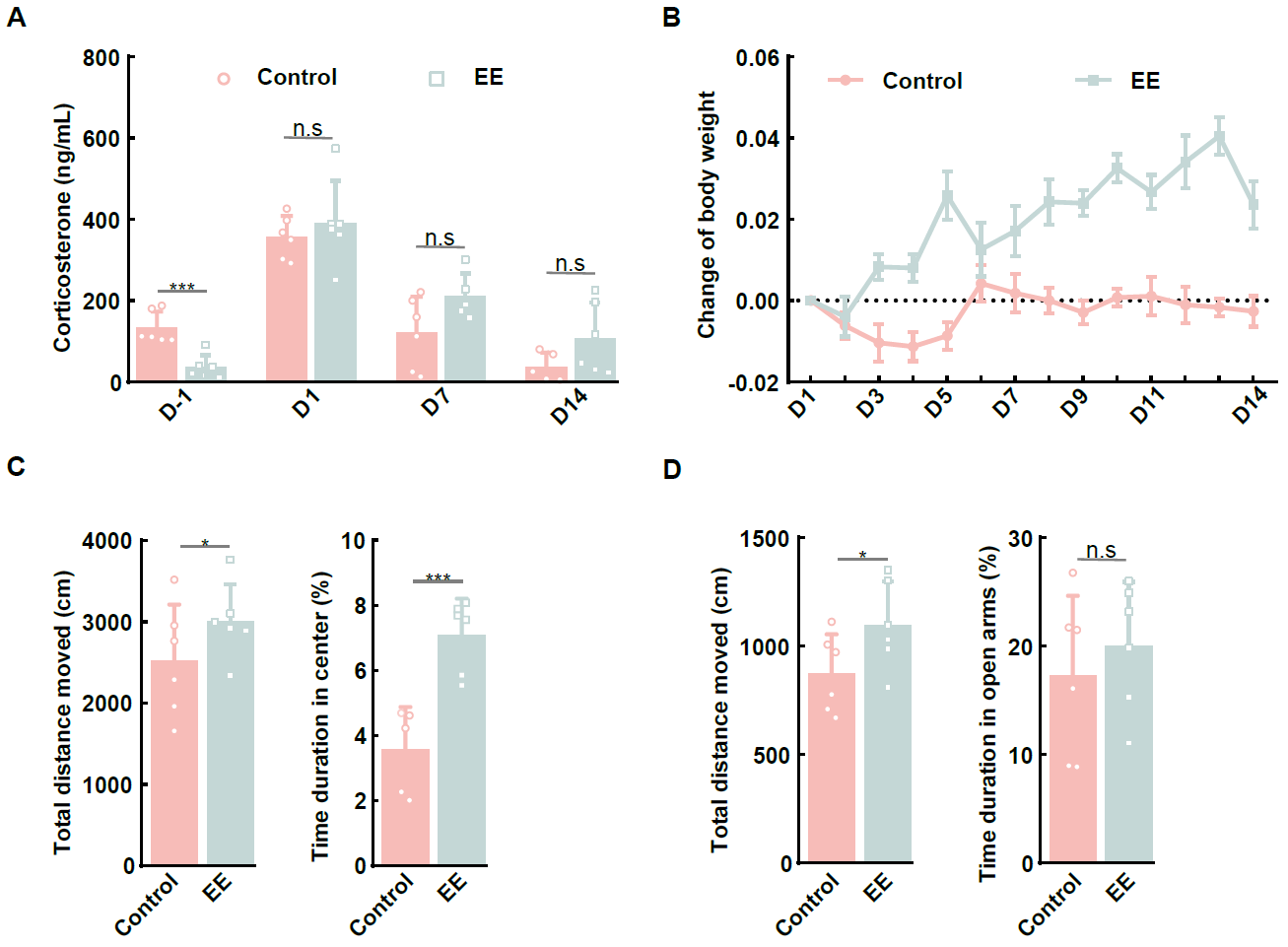

Figure S4. Effects of enriched environment (EE) on corticosterone level and behavior of awake mice secured with restraining setup 3 (RS3). (A) Comparison of corticosterone level in EE and control groups. (B) Change of body weight (mean ± SEM) in EE and control groups. (C, D) Open filed test (OFT) and elevated plus maze (EPM) results in EE and control groups. (n = 6, * p <0.05, *** p < 0.001)
