## Supplemental Figure 5 for "A systematically optimized awake mouse fMRI paradigm"

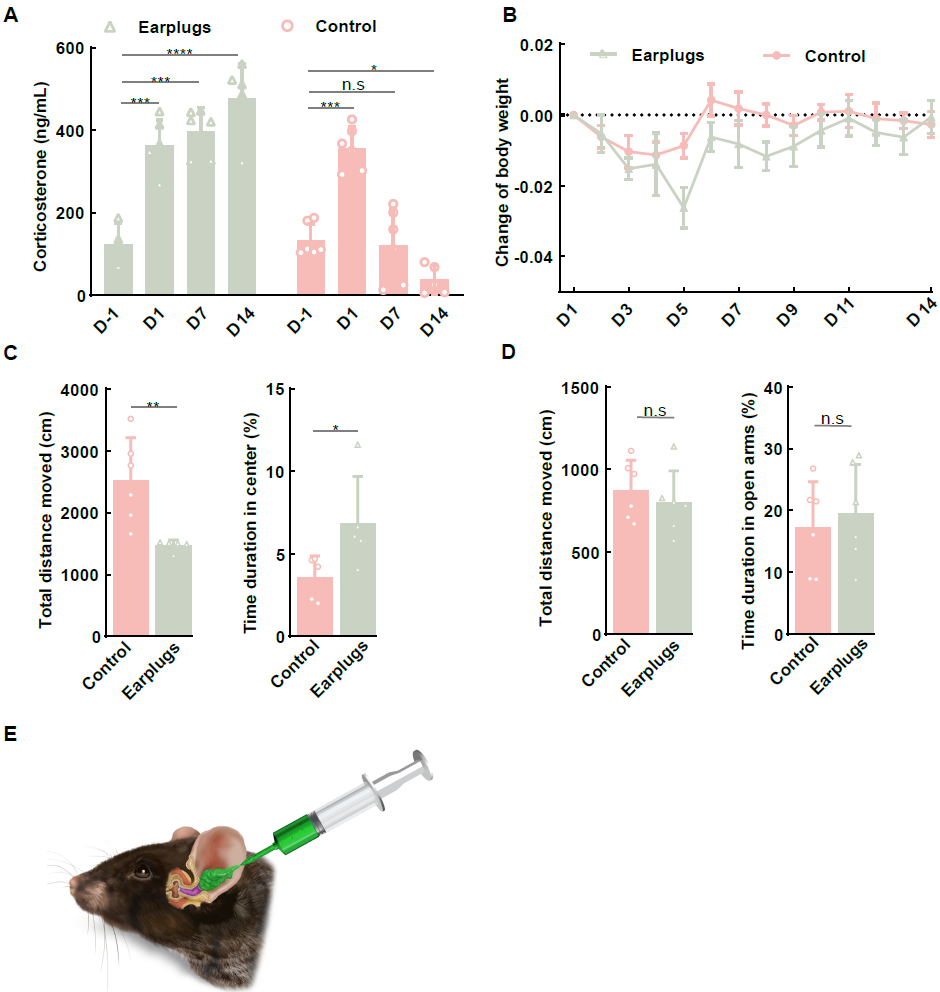

Figure S5. Effects of earplugs on corticosterone level and behavior of awake mice secured with RS3. (A) Corticosterone level in earplugs and control groups. (B) Change of body weight (mean ± SEM) during habituation in earplugs and control groups. (C, D) OFT and EPM results in earplugs and control groups. (E) The schematic of earplugs. (n = 6, *p < 0.05, **p <0.01, ***p < 0.001)
