## Supplemental Figure 6 for "A systematically optimized awake mouse fMRI paradigm"

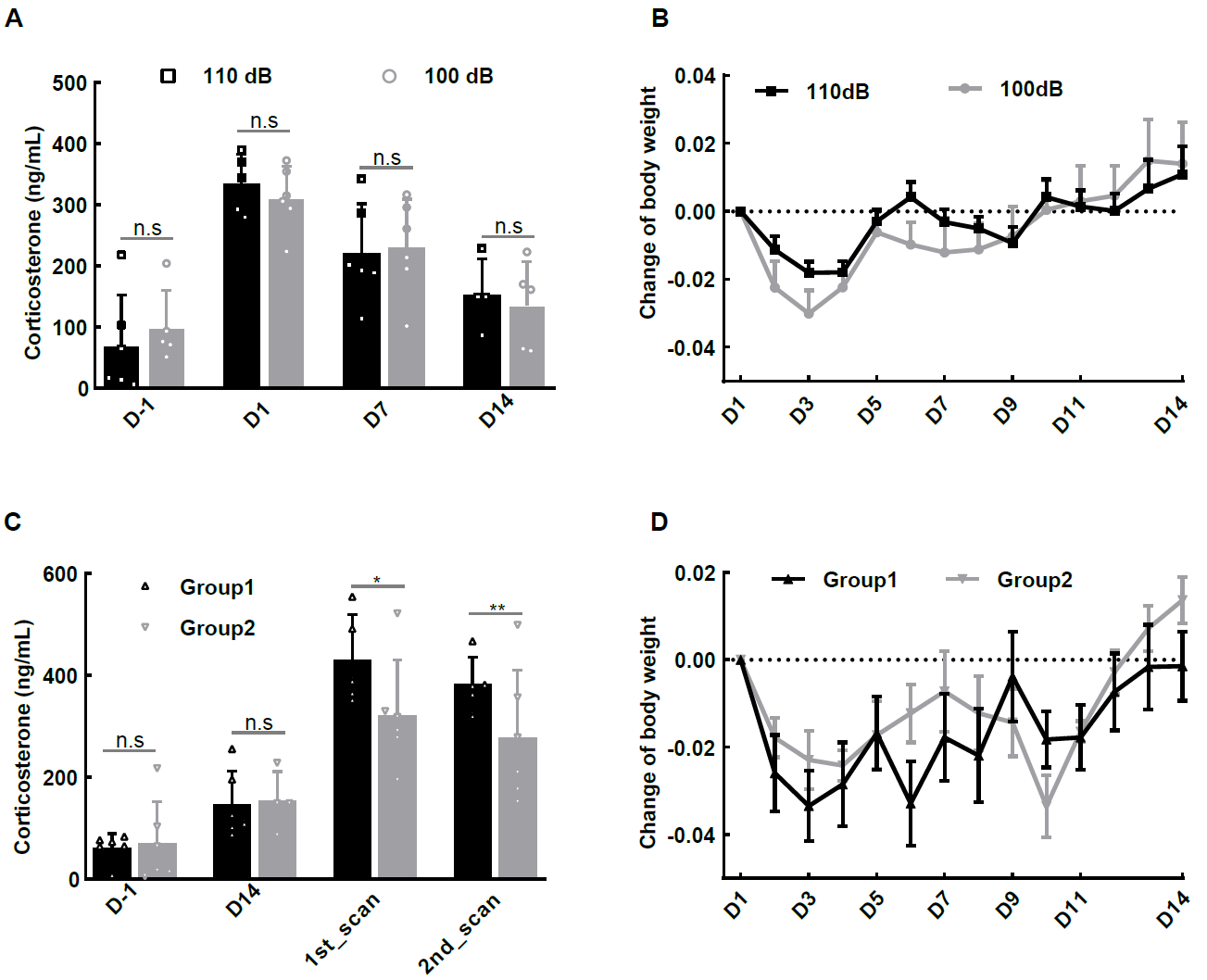

Figure S6. Effects of sound intensity on corticosterone level (A) and body weight. (B) in awake mice during habituation. Effects of MRI scanning duration on corticosterone level (C) and body weight (D) in awake mice. Group1: MRI scanning duration was 60 min; Group2: MRI scanning duration was 30 min. (n = 6, * p < 0.05, ** p < 0.01)
