## Supplemental Figure 7 for "A systematically optimized awake mouse fMRI paradigm"

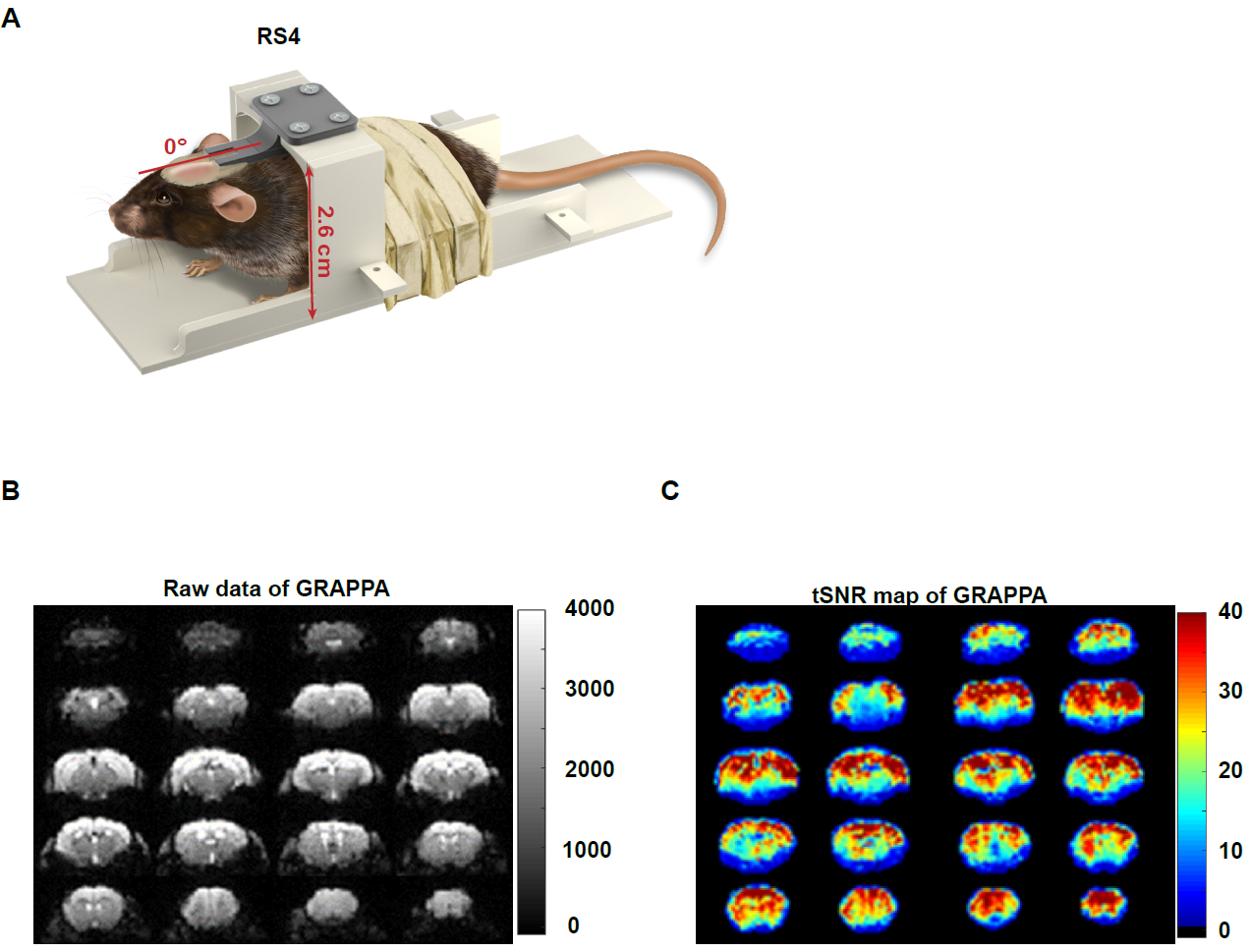

Figure S7. (A) The restraining setup (RS4) used with cryogenic mouse head coil (Bruker) for MRI data acquisition. (B) Raw GRAPPA EPI data. (C) Temporal signal-to-noise ratio (tSNR) map of GRAPPA-EPI.
