## Supplemental Figure 8 for "A systematically optimized awake mouse fMRI paradigm"

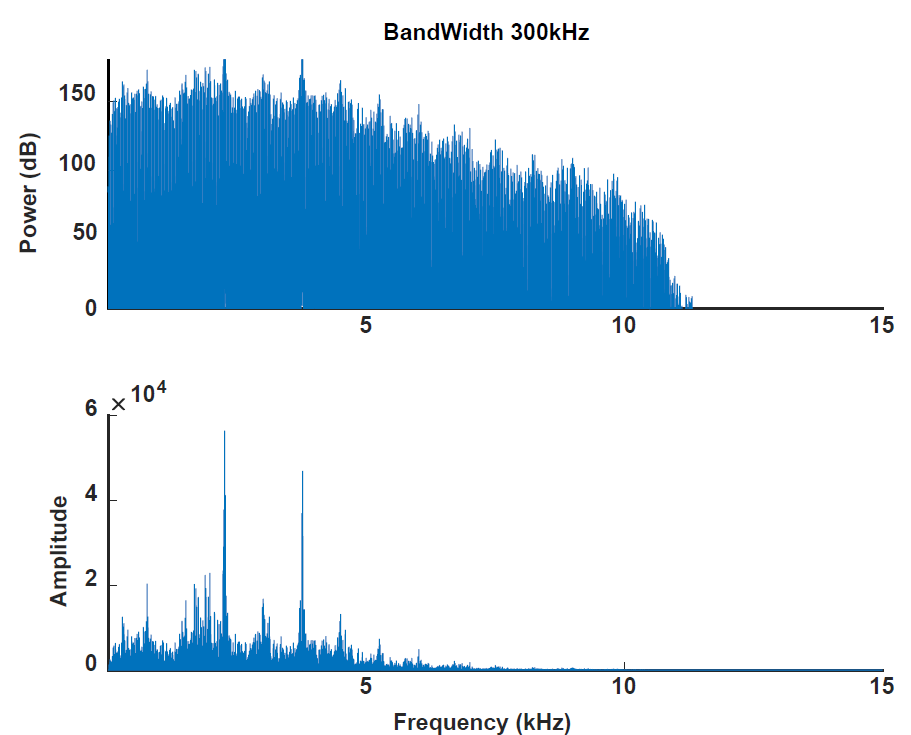

Figure S8. The fMRI-EPI noise recorded by fiber-optic microphone. The frequency of the fMRI-EPI noise is at relatively low frequency range, to which mice are insensitive.
