## Supplemental Table 1 for "A systematically optimized awake mouse fMRI paradigm"

Table S1. Summary of fMRI acquisition parameters (R: acceleration factor of iPAT; MBF: multiband factor).

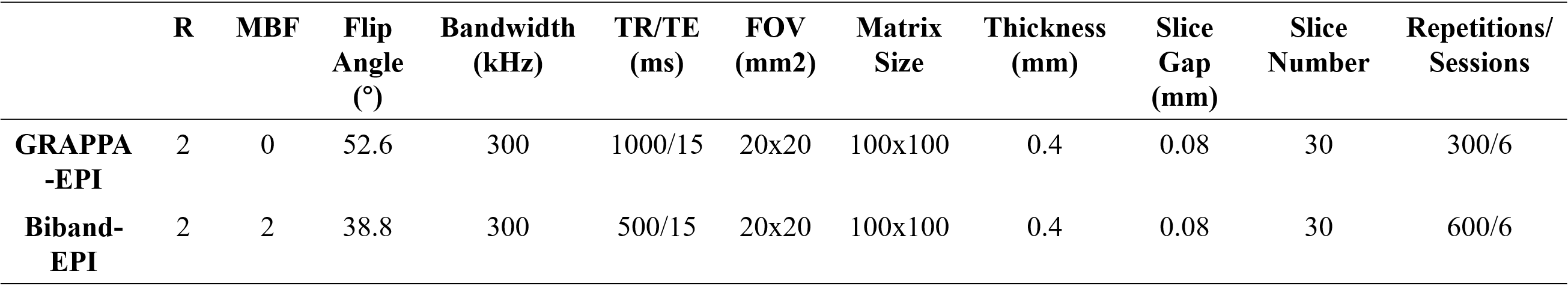
